# FAO-supported OxPhos leukemic stem cells are sensitive to cold

**DOI:** 10.1101/2023.04.17.537132

**Authors:** Emmanuel Griessinger, Diego Pereira-Martins, Marielle Nebout, Claudie Bosc, Estelle Saland, Emilie Boet, Ambrine Sahal, Johanna Chiche, Delphine Debayle, Lucille Fleuriot, Maurien Pruis, Véronique De Mas, François Vergez, Christian Récher, Gerwin Huls, Jean-Emmanuel Sarry, Jan Jacob Schuringa, Jean-François Peyron

## Abstract

Targeting mitochondrial oxidative phosphorylation (OxPhos) metabolism has revealed a potential weakness for leukemic stem cells (LSCs) that can be exploited for therapeutic purposes. Fatty acids oxidation (FAO) is a crucial OxPhos-fueling catabolic pathway for some AML and for chemotherapy-resistant AML cells. Here, we identified cold sensitivity at 4°C (cold killing challenge: CKC4), as a novel vulnerability that selectively kills FAO-supported OxPhos LSCs in Acute Myeloid Leukemia while sparing normal hematopoietic stem cells (HSCs). Cell death of OxPhos leukemic cells was induced by membrane permeabilization at 4°C while by sharp contrast, leukemic cells relying on glycolysis were resistant. Forcing glycolytic cells into OxPhos metabolism sensitized them to CKC4. We show using lipidomic and proteomic analyzes that OxPhos shapes the composition of the plasma membrane and introduce variation of 22 lipid subfamilies between cold-sensitive and cold-resistant cells. Cold sensitivity is a potential OxPhos biomarker.

**Significance:** This study reveals that mitochondrial energetics fueled by FAO metabolism introduces membrane fragility upon cold exposure in OxPhos-driven AMLs and in LSCs. This novel physical property of Leukemic cells and LSCs opens new avenues for biomarker and diagnostics as well as for anti-OxPhos drug screening and LSCs targeting.

**One Sentence Summary:** OxPhos leukemic cells die at 4°C

## Main Text

### Introduction

Cells rely on two main energetic pathways: glycolysis and oxidative phosphorylation (OxPhos), also called the mitochondrial metabolism that are fueled by amino acids, monosaccharides, and fatty acid (FA) pathway-derived substrates. OxPhos-driven metabolism predicts poor clinical outcomes for patients with certain types of solid tumors and leukemia (1–3). Metabolism-modulating agents, such as CPI-613 which prevents the entry of acetyl-CoA into the TCA cycle, or IACS-010759 which inhibits the mitochondria complex I, have shown promise in early-stage preclinical studies and clinical trials, either alone or in combination with other classes of agents (3–9). Here we uncover that steady-state energy metabolism at 37 degrees Celsius (37°C) predetermines the sensitivity of blasts and LSCs from AML patients subjected to a low temperature (4°C) commonly used as a storage procedure. While adaptive cell signaling has been widely described for normal stem cells in whole animal models or organs subjected to hypothermia, the survival of isolated cancer cells or subpopulations of cancer stem cells subjected to cold has not been detailed nor linked to OxPhos.

## Results

### Inter-patient and intra-patient cold sensitivity heterogeneity

Primary AML patient samples (Table S1) incubated in culture medium at 4°C significantly lose their viability over time as evidenced by an increased positivity to 4′,6-diamidino-2-phenylindole (DAPI) staining when the cells have regained their normothermia (Figure 1A, B). While some samples were highly sensitive to death induced by the cold killing challenge at 4°C (CKC4), others appeared resistant. Analysis of CKC4 responses revealed that the time to lose the first quartile of the viability of the population (Q1V^-^ Time) was an efficient early and simple parameter allowing to rank the cold sensitivity. A strong variation in Q1V^-^ Time between AML samples was observed, ranging from 4h to 300h with a median value of 96.5 SEM±7.14 hours. AML with a high Q1V^-^ Time had a significantly higher white blood count (WBC) (OR: 6.9; p=0.034) and harbored almost four times more the FMS-like tyrosine kinase-3 internal tandem duplication (*FLT3*-ITD) (OR: 3.75; p=0.034) or *NPM1* mutations (OR: 4.5; p=0.04) (Figure S1A). No difference in 4°C sensitivity was seen for patient age or risk groups or response to treatment. Similar heterogeneity in cold sensitivity was observed with a panel of 21 commonly used human myeloid or lymphoid leukemic cell lines (Figure S1B-C). We questioned whether LSCs would also be sensitive to 4°C. We monitored the respective sensitivity to cold exposure of leukemic subpopulations bearing the hematopoietic stem cell phenotype (HSC-like; CD34^+^CD38^-^ CD45RA^-^), the lymphoid-primed multipotent progenitors (LMPP-like; CD34^+^CD38^-^CD45RA^+^), the committed common myeloid progenitors (CMP-like; CD34^+^CD38^+^CD45RA^-^CD123^+^), the granulocyte-macrophage progenitors (GMP-like; CD34^+^CD38^+^CD45RA^+^CD123^+^) and the megakaryocyte-erythroid progenitors phenotype (MEP-like; CD34^+^CD38^+^CD45RA^-^CD123^-^) (10–13) (Figure 1C). Eight AML samples that were CD34-negative were excluded. We evidenced for 48 samples that HSC-like AML cells were more sensitive to CKC4 as compared to their normal counterpart measured in Cord Blood samples (CB) or compared to MEP-like, CMP-like, GMP-like, and LMPP-like AML cells (Figure 1D). The sensitivity to CKC4 of HSC-like AML cells was independent of the initial bulk viability after thawing at different measurement times (Figure 1E). We also noticed that the bulk sensitivity did not depend on the abundance of the HSC-like cells which suggests that the loss of viability results from the cumulative sensitivity of different populations. Thus, cold sensitivity reveals differences between samples and between sample subpopulations.

**Figure 1:**
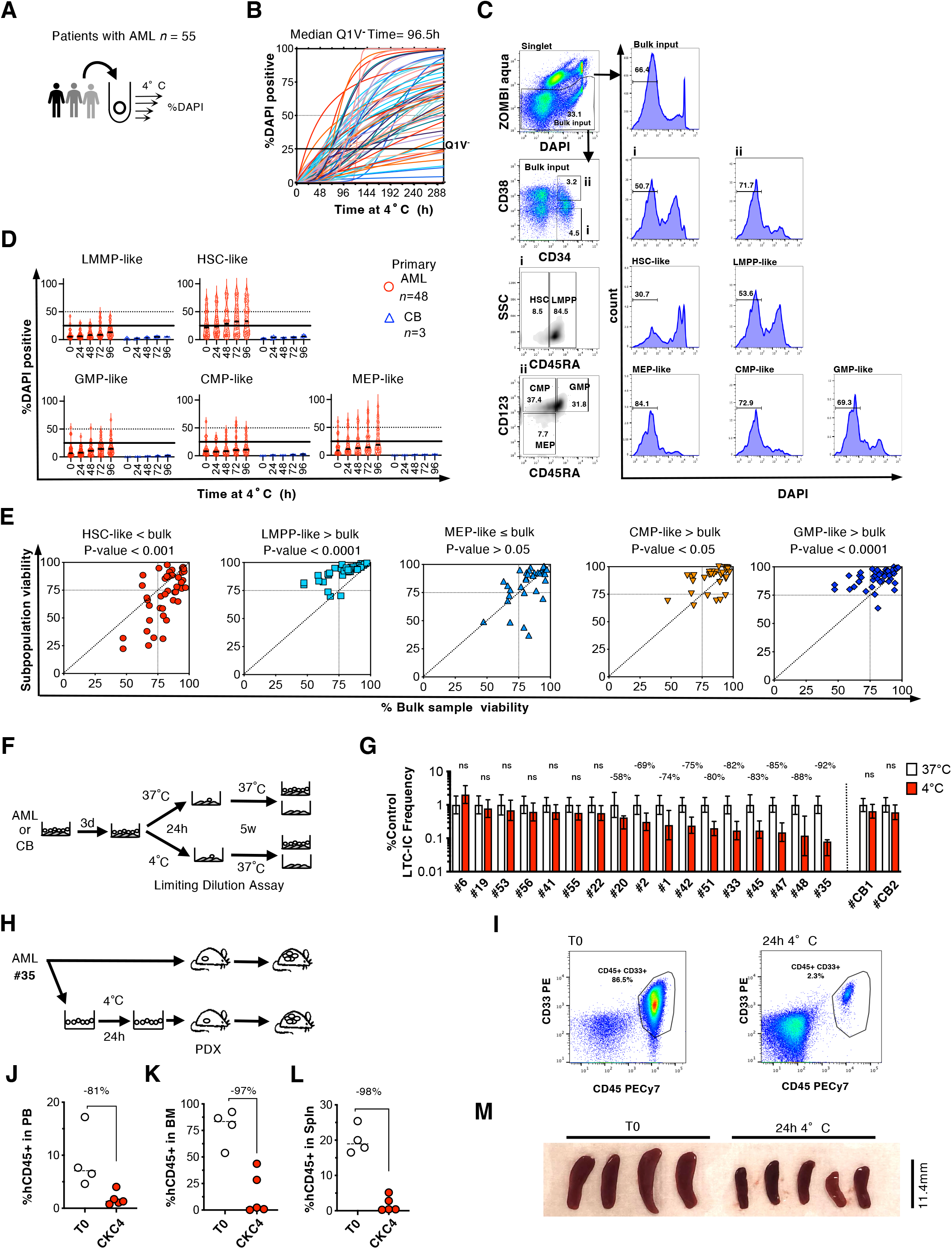
LSCs are sensitive to 4°C. (A) De novo primary AML samples (n=55, see Table S1) were incubated at 4°C and their viability were sequentially monitored over time by DAPI incorporation staining. (B) Kinetics of viability of primary bulk AML during the cold killing challenge at 4°C (CKC4). Median value of time to lose the first quartile of viability at 4°C (Q1V^-^ Time) determined for 55 AML samples. Curves shows the experimental derived non-linear regression trend line determined from 5 to 7 endpoint measurements. (C) FCM gating strategy. Input dead cells at thawing were excluded by Zombi-aqua negative selection to define the bulk leukemic population (Bulk input). Stem/progenitor-like (i: CD34^+^ CD38^-^) and progenitor-like (ii: CD34^+^ CD38^+^) were subdivided in Hematopoietic Stem Cells-like (HSC-like; CD34^+^CD38^-^CD45RA^-^), lymphoid-primed multipotential progenitors-like (LMPP-like; CD34^+^CD38^-^CD45RA^+^), committed common myeloid progenitors-like (CMP-like; CD34^+^CD38^+^CD45RA^-^CD123^+^), granulocyte-macrophage progenitors-like (GMP-like; CD34^+^CD38^+^CD45RA^+^CD123^+^) and megakaryocyte-erythroid progenitors-like (MEP-like; CD34^+^CD38^+^CD45RA^-^CD123^-^). The DAPI fluorescence profile was then monitored at different time for the Bulk input, the Stem/progenitor, the progenitor, the HSC, LMPP, CMP, GMP and the MEP-like gated populations. (D) Kinetics of viability of LMPP-like, HSC-like, GMP-like, CMP-like and MEP-like population subset within AML and CB samples during CKC4. (E) Subpopulations viabilities (Y-axis) against bulk population viability (X-axis) in primary AML samples. Data show the viability paired comparison determined for n=32 to 48 AML samples incubated at 4°C for 24h. Similar data were obtained at 48 and 72h. Comparisons were made using a paired Student’s t-test. (F) Experimental design for G. After 3 days of co-culture with MS-5, primary AML or CB CD34^+^ cells were either maintained at 37°C or incubated at 4°C for 24h then replated in limiting dilution analysis. Then the frequency of L-LTC-IC was determined after 5 weeks of culture at 37° C. (G) CKC4 impact on leukemic or normal LTC-IC. (H) Xenograft assay of primary AML cells undergoing or not a CKC4 for 24h. (I) Representative Flow cytometric analyses in mouse recipients of sample #35 from control condition (left) or CKC4 condition (right) are shown, indicating percentages of engrafted human AML cells (CD45^+^CD33^+^) in total bone marrow cells at 10 weeks. Quantifications of leukemic cells infiltration in PB (J), in the BM (K), and spleen (L) of AML #35. Each dot represents one transplanted mouse. (M) Spleen size. For F, I-K a Mann Whitney t-test was used for statistical analysis, the calculated percentage reductions are shown, bars represent median.

### LSC are sensitive to 4°C

LSCs can harbor membrane markers different from normal HSCs (13,14). Thus, we tested functionally how the CKC4 could impair the leukemic long-term culture initiating cells compartment (L-LTC-IC) that we have previously shown to be a faithful read-out for functionally defined LSC (15,16). We chose a 24h incubation period at 4°C, 4 times lower than the Q1V^-^ Time median of our cohort to fairly estimate the sensitivity of the LSCs. Seventeen AML samples and two CB samples were incubated either at 4°C or at 37°C for 24h, then frequencies of LTC-IC were determined at 37°C for both conditions (Figure 1F). No significant count differences were seen at the bulk population level after 24h at 4°C for AML and CB. However, for 10 AML samples out of the 17 tested (58%) a reduction in the frequency and the absolute number of L-LTC-IC was observed after CKC4 ranging from 58% to 92% reductions as compared to the constant 37°C experimental arm (Figure 1G). No 4°C impact was seen for the CB samples tested. We tested the CKC4 impact at the level of the SCID-Leukemia-Initiating Cells (SL-IC) through the xenograft assay (Figure 1H). Since in the in LTC experiments the control experimental arm maintain at 37°C arm may have benefit from a transient expansion we decided for SL-IC measurement to inject an equal number of live cells at T0 or 24h after the CKC4. Ten weeks later we observed a massive reduction of patient leukemia infiltration in the mice peripheral blood (PB), in their bone marrow (BM) and spleen (Spln), as well as a reduced splenomegaly in the experimental arms of the CKC4 (Figure 1I-M). This clearly shows that leukemic stem cells are selectively highly sensitive to cold exposure.

### OxPhos Leukemic Cells Exhibit a specific membranes permeabilization at 4°C

We noticed that CKC4 sensitivity did not correlate with freeze/thaw sensitivity, cell size, or cell division rate (Figure S1D-G). However, we evidenced that cold sensitivity depends on mitochondrial parameters measured at steady-state in normothermia. We measured a significant negative correlation between the Q1V^-^ Time parameter and the maximal and spare respiration capacity, the ATP-linked respiration, the mitochondrial DNA content (mtDNAc), the mitochondrial membrane potential, the calcium content as well as the amount of ATP generated by OxPhos (Figure 2A and Figure S1H-J). CKC4 responses were highly reproducible (Figure S1K). Cell death at 4°C was characterized by permeabilization of the plasma membrane and the organelles membrane as evidenced by the leakage of fluorescent molecules DAPI, TMRE or GFP which occurred with a faster kinetic for OxPhos Cells (Figure S1L). The cell-specific membranes permeabilization at 4°C was confirmed by either performing CKC4 on cell lines that were biobanked and maintained in different laboratories or by testing the same cells in different laboratories, resulting in similar sensitivity profiles (Figure S1M-N). Altogether, our results suggest that leukemic cells relying on OxPhos metabolism are specifically sensitive to CKC4.

**Figure 2.**
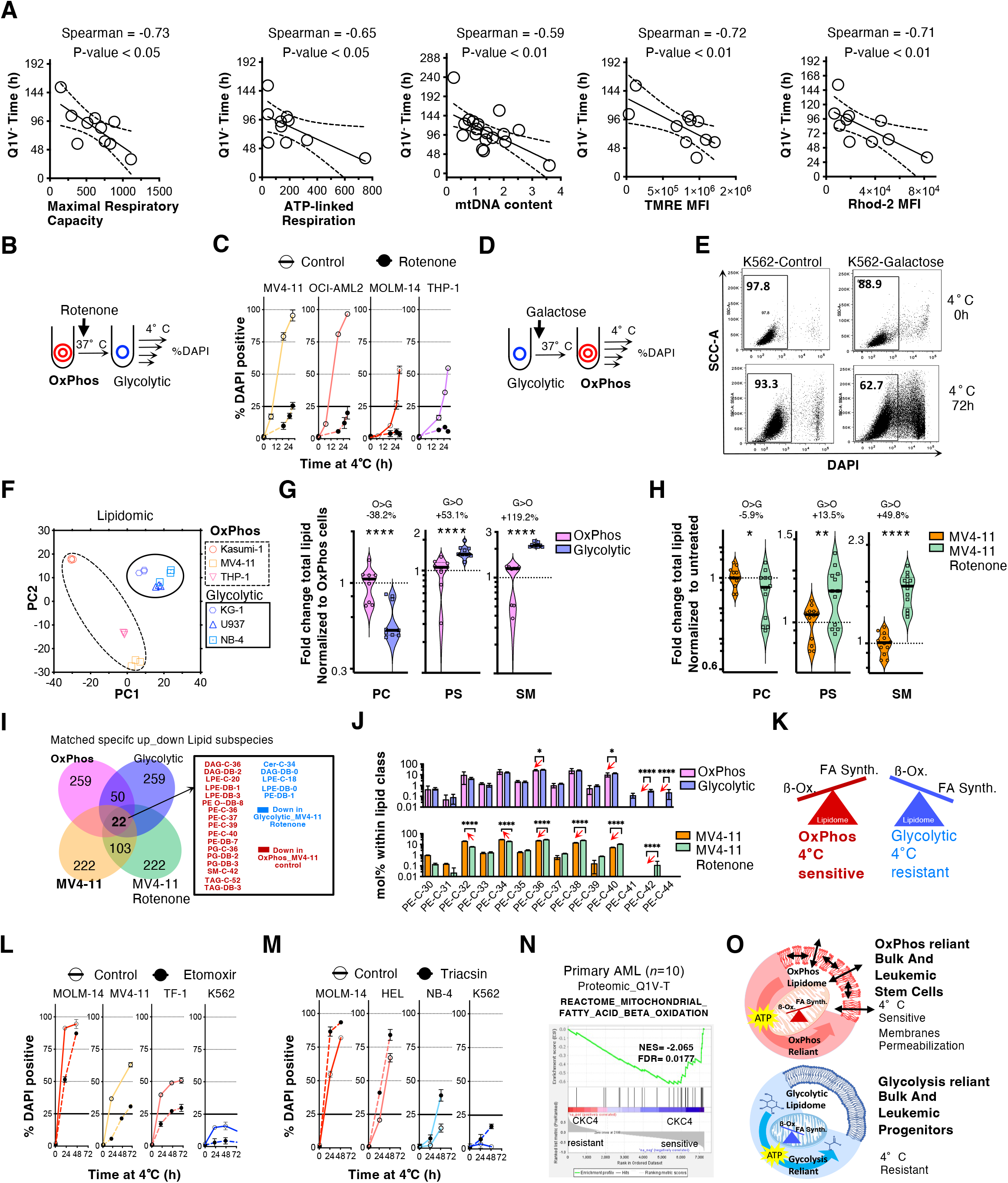
OxPhos leukemic cells have a 4°C sensitive specific lipidome dependent on fatty acid metabolism. (A) Maximal Respiratory Capacity, ATP-linked respiration, mitochondrial DNA content (mtDNAc), median fluorescence intensity (MFI) for the mitochondrial potential probe TMRE, and the mitochondrial calcium probe Rhod-2 as a function of Q1V^-^ Time at 4°C for primary AML. Respiration 5 was measured with Seahorse assays under basal and maximal conditions. The relative mtDNA copy number was calculated relative to a reference DNA sample (healthy donor) set to 1. Non-parametric Spearman correlation tests were applied for n=10 to 18 pairs. Thin black line shows experimental derived linear regression trend line with 95% confidence band (B) Glycolytic-forced cells submitted to CKC4. Mitochondrial respiration was impaired by the complex I inhibitor rotenone. (C) Kinetic of 10 Viability of Pre-rotenone or untreated control AML cells at 4°C. (D) OxPhos-forced cells submitted to CKC4. K562 cells were incubated at 37°C in glucose-free medium with galactose for 72h. (E) Representative FACS dot plot of side scatter against DAPI for untreated control or Galactose pre-treated K562 cells after 72h at 4°C. (F) PCA score plot of the total lipidome by MS (including species and subspecies analysis) of three OxPhos (Kasumi-1, THP-1, MV4-11) and three glycolytic leukemic 15 cells (KG-1a, NB-4, U937), triplicate per cell type. (G) Fold change of ether-phosphatidylcholine (PC), Phosphatidylserine (PS) and Sphingomyelin (SM) for OxPhos (O) and glycolytic cells (G) indicated in G normalized to the mean percentage for OxPhos cells (see also Table 1 and Fig. S2I). (H) Fold change of PC, PS and SM for untreated OxPhos MV4-11 and glycolytic reprogrammed Rotenone treated MV4-11 cells. A Mann–Whitney test was applied, *p<0.05, **p < 0.01, ****p < 0.0001. (I) Venn 20 diagram of significantly different proportion of lipid subspecies for OxPhos and glycolytic cells cohorts and for MV4-11 and MV4-11 rotenone (see also Table 2). (J) Repartition of PE-C-30 to PE-C-44 subspecies within the PE family for OxPhos and glycolytic cells cohorts and for MV4-11 and MV4-11 rotenone treated cells. (K) Diagram showing the hypothesis tested in M-N. (L) Indicated cells were pretreated with non-toxic dose of mitochondrial Beta-oxidation inhibitor Etomoxir or (M) with lipid 25 synthesis inhibitor triacsin C for 72h before undergoing a CKC4. (N) Total proteome by MS for 10 patient samples. Pearson correlations were calculated for all quantified proteins versus Q1V^-^ Time determined, and a ranked list of Pearson coefficients was used to perform gene set enrichment analyses (see also Table S2). GSEA analysis of Fatty Acid Beta Oxidation and the HSC up pathways in CKC4 resistant versus CKC4 sensitive primary AML samples. (O) Overview of study findings. 30 OxPhos reliant blast and LSCs cells have a FA specific metabolism shaping the composition of their membranes and accounting for their faster permeabilization compared to glycolysis-dependent leukemic blasts and progenitors.

### The metabolic state shapes the cellular cold sensitivity

Since enzymatic activities are impeded at 4°C, we hypothesized that the metabolism at 37°C predetermines the sensitivity to CKC4. We wondered whether we could modify the CKC4 response by directly influencing the metabolic state at 37°C prior to cold exposure. We identified different OxPhos-dependent cell lines that tolerated the blockade of the mitochondrial complex I by rotenone and were consequently driven into glycolysis, as evidenced by an increased lactate secretion (MOLM-14, MV4-11, OCI-AML2, THP-1; Figure 2B and Figure S2A). These glycolytic-forced cells became CKC4 resistant (Figure 2C). This induced resistance required at least 24 hours of metabolic reprogramming at 37°C (Figure S2B). We next undertook the opposite experiment by forcing glycolytic cells to become OxPhos (Figure 2D). K562 cells incubated for 72h with the glucose stereoisomer galactose could switch their energy production towards OxPhos (Figure S2C) (17). Control K562 cells were cold resistant, but became sensitive upon pretreatment with galactose (Figure 2E). NB4 cells can be differentiated upon treatment with all-trans retinoic acid (ATRA), coinciding with decreased glycolysis and increased in OxPhos (Figure S2D-E). ATRA-treated NB4 became CKC4 sensitive (Figure S2F). And similar to what was observed during the metabolic reprogramming of OxPhos cells towards glycolysis, the OxPhos switch of NB4 cells also required a long period to influence the CKC4 response (Figure S2G), suggesting that sensitivity to cold exposure reflects a stable and pre-established mitochondria-dependent metabolic profile. Overall, modulating OxPhos influenced the cellular cold sensitivity which reveals a causal relationship between the steady-state cellular metabolism and the cell death at 4°C.

**Table 1:** OxPhos and Glycolytic AML cells Lipid Classes percentages. This table contains the percentage of amount of a lipid class calculated by summing the pmol values of the individual lipids belonging to each class. Class amount was then normalized to total lipid content. p-value, Mann-Whitney t-test was applied to compare OxPhos (O: Kasumi-1, THP-1, MV4-11) and glycolytic cells (G: KG-1, NB-4, U937), ns: not significant. n=3 per cell line. CE: Cholesterol esters. CER: Ceramide. CL: Cardiolipin. DAG: Diacylglycerol. HEXCER: Hexosylceramide. LPC: lyso-Phosphatidylcholine. LPC: (O-) lyso-Phosphatidylcholine (-ether). LPE: lyso-Phosphatidylethanolamine. LPE: (O-) lyso-Phosphatidylethanolamine (-ether). PA: Phosphatidate. PC: Phosphatidylcholine. PC (O-): Phosphatidylcholine (-ether). PE: Phosphatidylethanolamine. PE (O-): Phosphatidylethanolamine (-ether). PG: Phosphatidylglycerol. PI: Phosphatidylinositol. PS: Phosphatidylserine. SM: Sphingomyelin. TAG: Triacylglycerol.

|  | Kazumi-1<br>(O) | THP-1 (O) | MV4-11 (O) | KG-1 (G) | NB-4 (G) | U937 (G) | Percentage<br>change<br>(O-G) | Fold<br>percentage<br>change<br>(O/G) | P-value |
| --- | --- | --- | --- | --- | --- | --- | --- | --- | --- |
| **CE** | 0,456 ± 0,608044 | 0,487 ± 0,032 | 0,1158 ± 0,154461 | 0,821 ± 0,000 | 0,964 ± 0,964 | 0,174 ± 0,000 | ns | ns | 5,85E-01 |
| **Cer** | 0,398 ± 0,009 | 0,617 ± 0,011 | 0,294 ± 0,028 | 0,108 ± 0,013 | 0,097 ± 0,008 | 0,096 ± 0,003 | **0,336** | **4,340** | 1,09E-04 |
| **CL** | 0,439 ± 0,127 | 0,734 ± 0,074 | 1,226 ± 0,633 | 1,017 ± 0,102 | 0,704 ± 0,096 | 1,019 ± 0,048 | ns | ns | 5,80E-01 |
| **DAG** | 2,725 ± 0,061 | 1,783 ± 0,097 | 1,080 ± 0,123 | 0,570 ± 0,057 | 0,543 ± 0,024 | 0,840 ± 0,044 | **1,211** | **2,861** | 9,20E-04 |
| **HexCer** | 0,018 ± 0,003 | 0,319 ± 0,012 | 0,651 ± 0,110 | 0,115 ± 0,023 | 0,136 ± 0,021 | 0,210 ± 0,010 | ns | ns | 1,03E-01 |
| **LPC** | 0,024 ± 0,001 | 0,213 ± 0,008 | 0,086 ± 0,011 | 0,048 ± 0,002 | 0,057 ± 0,007 | 0,047 ± 0,001 | ns | ns | 7,55E-02 |
| **LPC O-** | 0 ± 0 | 0,113 ± 0,007 | 0,036 ± 0,024 | 0,016 ± 0,000 | 0,007 ± 0,007 | 0,021 ± 0,006 | **0,072** | **5,045** | 8,02E-03 |
| **LPE** | 0,022 ± 0,001 | 0,117 ± 0,002 | 0,061 ± 0,005 | 0,052 ± 0,008 | 0,063 ± 0,006 | 0,049 ± 0,003 | ns | ns | 4,00E-01 |
| **LPE O-** | 0,007 ± 0,000 | 0,036 ± 0,001 | 0,019 ± 0,002 | 0,005 ± 0,000 | 0,003 ± 0,002 | 0,004 ± 0,000 | **0,016** | **4,733** | 4,89E-03 |
| **PA** | 0 ± 0 | 1,317 ± 0,082 | 0,399 ± 0,241 | 0,1123 ± 0,075 | 0,556 ± 0,042 | 0,393 ± 0,058 | ns | ns | 9,47E-02 |
| **PC** | 37,05 ± 1,046 | 42,70 ± 0,785 | 41,78 ± 5,067 | 43,82 ± 1,375 | 44,78 ± 0,684 | 43,87 ± 0,292 | ns | ns | 4,12E-02 |
| **PC O-** | 4,631 ± 0,087 | 6,747 ± 0,159 | 7,117 ± 0,844 | 3,180 ± 0,081 | 3,098 ± 0,112 | 5,140 ± 0,127 | **2,359** | **1,620** | 6,35E-04 |
| **PE** | 11,15 ± 0,206 | 10,49 ± 0,132 | 14,05 ± 0,399 | 15,89 ± 0,311 | 16,86 ± 0,397 | 14,24 ± 0,144 | **-3,768** | **0,759** | 7,25E-05 |
| **PE O-** | 8,831 ± 0,159 | 11,71 ± 0,147 | 15,09 ± 0,148 | 8,884 ± 0,187 | 7,162 ± 0,021 | 8,071 ± 0,238 | **3,837** | **1,477** | 2,60E-03 |
| **PG** | 0,649 ± 0,047 | 0,661 ± 0,023 | 0,819 ± 0,054 | 0,092 ± 0,013 | 0,168 ± 0,014 | 0,191 ± 0,005 | **0,559** | **4,717** | 4,91E-09 |
| **PI** | 8,074 ± 0,734 | 7,682 ± 0,663 | 7,253 ± 3,496 | 9,713 ± 0,523 | 9,408 ± 0,534 | 10,14 ± 0,935 | ns | ns | 4,16E-02 |
| **PS** | 5,474 ± 0,773 | 6,928 ± 0,244 | 5,386 ± 2,384 | 9,395 ± 1,511 | 8,893 ± 0,661 | 8,944 ± 0,539 | **-3,148** | **0,653** | 8,97E-04 |
| **SM** | 3,618 ± 0,069 | 3,538 ± 0,067 | 1,322 ± 0,173 | 6,252 ± 0,259 | 6,336 ± 0,336 | 5,995 ± 0,098 | **-3,368** | **0,456** | 1,02E-05 |
| **TAG** | 16,44 ± 0,298 | 3,798 ± 0,095 | 3,220 ± 0,477 | 0,463 ± 0,100 | 0,490 ± 0,049 | 0,669 ± 0,148 | **7,279** | **14,468** | 9,80E-03 |

**Table 2:**
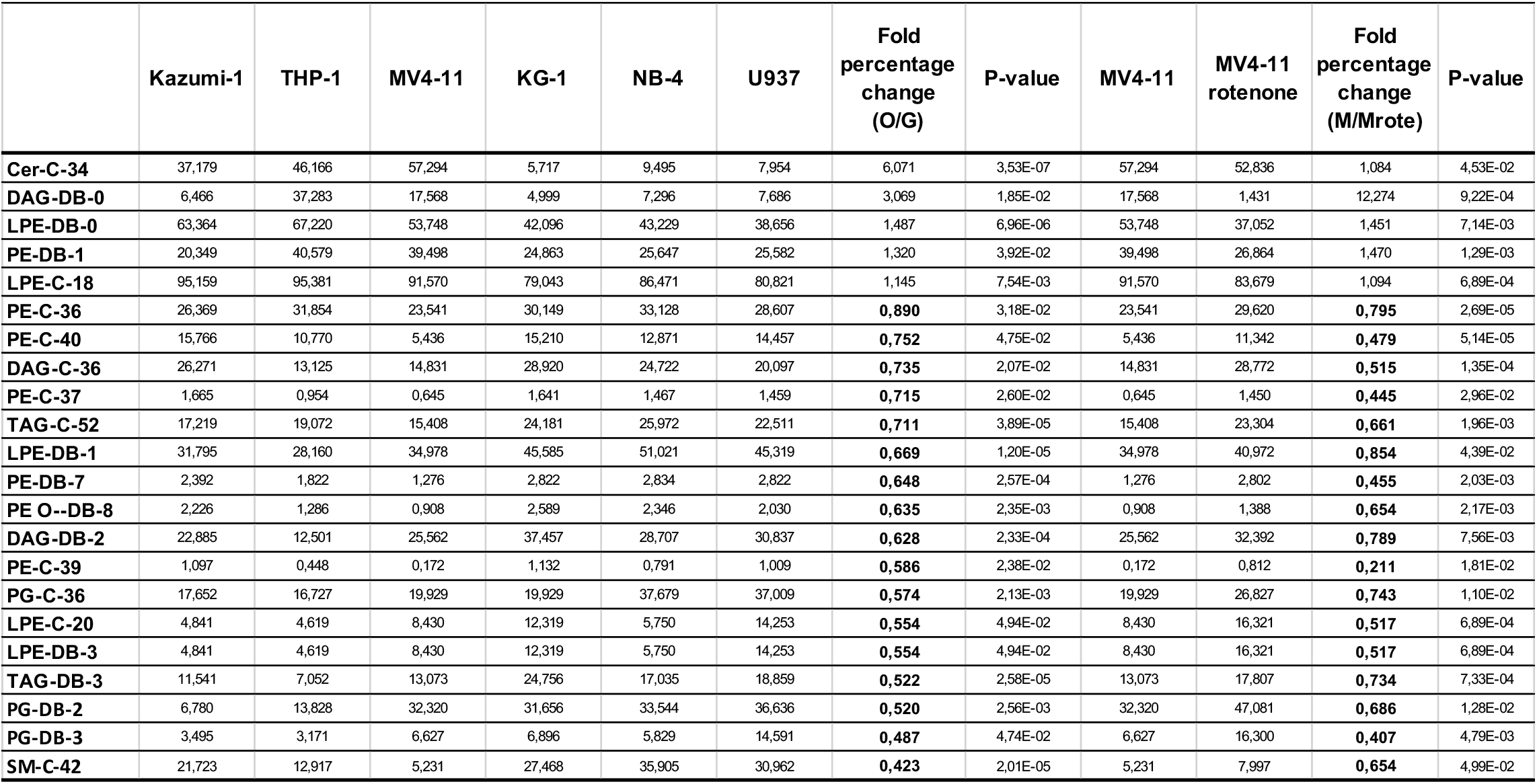
Lipid subspecies enriched or reduced between OxPhos versus glycolytic and between MV4-11 versus glycolytic-forced MV4-11 AML cells. Lipid subclass amount percentages was normalized with the summing the pmol values of the individual lipids belonging to each subclass. p-value, Mann-Whitney t-test was applied to compare OxPhos (O: Kasumi-1, THP-1, MV4-11) and glycolytic cells (G: KG-1, NB-4, U937) and MV4-11 (M) versus glycolytic-forced rotenone treated MV4-11 cells (Mrote). Lipid subclasses were ranked by the fold percentage change measured for O/G.

|  | Kazumi-1 | THP-1 | MV4-11 | KG-1 | NB-4 | U937 | Fold percentage change (O/G) | P-value | MV4-11 | MV4-11 rotenone | Fold percentage change (M/Mrote) | P-value |
| --- | --- | --- | --- | --- | --- | --- | --- | --- | --- | --- | --- | --- |
| **Cer-C-34** | 37,179 | 46,166 | 57,294 | 5,717 | 9,495 | 7,954 | 6,071 | 3,53E-07 | 57,294 | 52,836 | 1,084 | 4,53E-02 |
| **DAG-DB-0** | 6,466 | 37,283 | 17,568 | 4,999 | 7,296 | 7,686 | 3,069 | 1,85E-02 | 17,568 | 1,431 | 12,274 | 9,22E-04 |
| **LPE-DB-0** | 63,364 | 67,220 | 53,748 | 42,096 | 43,229 | 38,656 | 1,487 | 6,96E-06 | 53,748 | 37,052 | 1,451 | 7,14E-03 |
| **PE-DB-1** | 20,349 | 40,579 | 39,498 | 24,863 | 25,647 | 25,582 | 1,320 | 3,92E-02 | 39,498 | 26,864 | 1,470 | 1,29E-03 |
| **LPE-C-18** | 95,159 | 95,381 | 91,570 | 79,043 | 86,471 | 80,821 | 1,145 | 7,54E-03 | 91,570 | 83,679 | 1,094 | 6,89E-04 |
| **PE-C-36** | 26,369 | 31,854 | 23,541 | 30,149 | 33,128 | 28,607 | **0,890** | 3,18E-02 | 23,541 | 29,620 | **0,795** | 2,69E-05 |
| **PE-C-40** | 15,766 | 10,770 | 5,436 | 15,210 | 12,871 | 14,457 | **0,752** | 4,75E-02 | 5,436 | 11,342 | **0,479** | 5,14E-05 |
| **DAG-C-36** | 26,271 | 13,125 | 14,831 | 28,920 | 24,722 | 20,097 | **0,735** | 2,07E-02 | 14,831 | 28,772 | **0,515** | 1,35E-04 |
| **PE-C-37** | 1,665 | 0,954 | 0,645 | 1,641 | 1,467 | 1,459 | **0,715** | 2,60E-02 | 0,645 | 1,450 | **0,445** | 2,96E-02 |
| **TAG-C-52** | 17,219 | 19,072 | 15,408 | 24,181 | 25,972 | 22,511 | **0,711** | 3,89E-05 | 15,408 | 23,304 | **0,661** | 1,96E-03 |
| **LPE-DB-1** | 31,795 | 28,160 | 34,978 | 45,585 | 51,021 | 45,319 | **0,669** | 1,20E-05 | 34,978 | 40,972 | **0,854** | 4,39E-02 |
| **PE-DB-7** | 2,392 | 1,822 | 1,276 | 2,822 | 2,834 | 2,822 | **0,648** | 2,57E-04 | 1,276 | 2,802 | **0,455** | 2,03E-03 |
| **PE O--DB-8** | 2,226 | 1,286 | 0,908 | 2,589 | 2,346 | 2,030 | **0,635** | 2,35E-03 | 0,908 | 1,388 | **0,654** | 2,17E-03 |
| **DAG-DB-2** | 22,885 | 12,501 | 25,562 | 37,457 | 28,707 | 30,837 | **0,628** | 2,33E-04 | 25,562 | 32,392 | **0,789** | 7,56E-03 |
| **PE-C-39** | 1,097 | 0,448 | 0,172 | 1,132 | 0,791 | 1,009 | **0,586** | 2,38E-02 | 0,172 | 0,812 | **0,211** | 1,81E-02 |
| **PG-C-36** | 17,652 | 16,727 | 19,929 | 19,929 | 37,679 | 37,009 | **0,574** | 2,13E-03 | 19,929 | 26,827 | **0,743** | 1,10E-02 |
| **LPE-C-20** | 4,841 | 4,619 | 8,430 | 12,319 | 5,750 | 14,253 | **0,554** | 4,94E-02 | 8,430 | 16,321 | **0,517** | 6,89E-04 |
| **LPE-DB-3** | 4,841 | 4,619 | 8,430 | 12,319 | 5,750 | 14,253 | **0,554** | 4,94E-02 | 8,430 | 16,321 | **0,517** | 6,89E-04 |
| **TAG-DB-3** | 11,541 | 7,052 | 13,073 | 24,756 | 17,035 | 18,859 | **0,522** | 2,58E-05 | 13,073 | 17,807 | **0,734** | 7,33E-04 |
| **PG-DB-2** | 6,780 | 13,828 | 32,320 | 31,656 | 33,544 | 36,636 | **0,520** | 2,56E-03 | 32,320 | 47,081 | **0,686** | 1,28E-02 |
| **PG-DB-3** | 3,495 | 3,171 | 6,627 | 6,896 | 5,829 | 14,591 | **0,487** | 4,74E-02 | 6,627 | 16,300 | **0,407** | 4,79E-03 |
| **SM-C-42** | 21,723 | 12,917 | 5,231 | 27,468 | 35,905 | 30,962 | **0,423** | 2,01E-05 | 5,231 | 7,997 | **0,654** | 4,99E-02 |

### OxPhos leukemic cells have a CKC4 sensitive specific lipidome dependent on fatty acid metabolism

We then investigated which molecular mechanisms could underlie the specific sensitivity of OxPhos cells during CKC4. We performed a Pearson correlation analysis using the Q1V^-^ Time and a metabolomics dataset from the Cancer Cell Line Encyclopedia (CCLE) (18) which uncovered that many lipids had a clear abundance difference between OxPhos and glycolytic cells. To confirm this observation, we performed a complete lipidomic analysis of three OxPhos and three glycolytic AML cell lines. The principal component analysis of 1372 lipid subspecies confirmed a marked difference between the lipidomic profile of OxPhos and glycolytic leukemic cells (Figure 2F). 11 of the 19 classes of lipids, including energy storage and membrane lipids, were found different between OxPhos and glycolytic cells. OxPhos cells had a higher proportion of ether-phosphatidylethanolamine (PE O-), ether-phosphatidylcholine (PC), Triacylglycerol (TAG), Diacylglycerol (DAG), Phosphatidylglycerol (PG), Ceramide (CER), ether-lyso-Phosphatidylcholine (LPC O-), ether-lyso-Phosphatidylethanolamine (LPE O-) and a lower proportion of PE, Phosphatidylserine (PS) and Sphingomyelin (SM) (Table 1, Figure 2G and Figure S2I). We also determined the lipidome of OxPhos MV4-11 cells treated with rotenone for 72h and detected a profound lipidome remodeling upon OxPhos interference mimicking certain aspects of the glycolytic cell lipidome profile such as a lower PC and higher PS and SM content (Figure 2H). 22 lipid subspecies were commonly enriched or reduced between OxPhos versus glycolytic cells and between MV4-11 versus glycolytic-forced MV4-11 cells. Notably, 77% of these lipid subspecies were found reduced in OxPhos cells (Table 2 and Figure 2I). Among the reduced lipids, we observed several PE subspecies (PE-C-36 to PE-C-40). PE was the second richest family of lipids representing on average 23.73%±2% of the total cellular lipids (Table 1). Among the PE family, the distribution of carbon chains of different lengths revealed that cells relying on the OxPhos metabolism have a shift towards shorter PE subspecies with an even number of carbons (Figure 2J). This suggested that OxPhos leukemic cells could have higher catabolism and/or a lower anabolism of some of their fatty acids (FA) through the 2-Carbon acetyl Co-A moiety via the FA beta-oxidation or the FA synthesis pathways, respectively, which could directly influence CKC4 response (Figure 2K). We tested this hypothesis by treating leukemic cells with nontoxic doses of etomoxir, a FA beta-oxidation inhibitor that targets the mitochondrial transporter CPT-1, and Triacsin C, an inhibitor of long fatty acyl CoA synthetase which interferes with FA synthesis. Consistently with our hypothesis, etomoxir reduced the CKC4 sensitivity while Triacsin C increased it either for OxPhos or glycolytic leukemic cells (Figure 2L-M). To extend these results to primary AML, we established a quantitative proteome for 10 patient samples (Hogeling et al, in progress). Pearson correlations were calculated for all quantified proteins versus Q1V^-^ Time (Table S2), and a ranked list of Pearson coefficients was used to perform gene set enrichment analyses (GSEA). Consistently with our prior functional assays, this analysis identified stem cell signatures in CKC4-sensitive AML patient samples (Figure S2J). However, since our proteomics dataset was performed on a population of CD34+ blasts, it is also possible that these signatures result from distinguishing between patient samples enriched in progenitors and samples enriched in more committed cells. These GSEA analyzes also revealed that CKC4-sensitive AML patient samples were significantly enriched for the terms mitochondrial and fatty acid beta-oxidation (Figure 2N), but also mitochondrial translation, pyruvate metabolism and citric acid cycle, respiratory electron transport, and oxidative phosphorylation pathways (FigureS2J) which confirmed the distinct mitochondrial activity measured above through functional analysis. Overall, our results show that cellular cold sensitivity is linked to a FA beta-oxidation-supported OxPhos that shapes the membrane lipidome. We also reveal for the first time that cold is deleterious for L-LTC-ICs and SL-ICs (Figure 2O).

## Discussion

In this study, we show that leukemic cells have a distinct sensitivity to cell death induced by cold exposure and that this sensitivity is shaped by their oxidative metabolism and settled in their membrane lipid composition. Our discovery that metabolism influences the membrane composition is supported by a prior report showing changes in glycerophospholipid acyl chain lengths in yeast strains in which different kinases and phosphatases involved in energetic metabolism pathways were knocked out (19). Different studies highlighted the importance of fatty acid metabolism and the cross-talk between LSCs and adipose tissue in the AML niche. LSCs overexpress the fatty acid transporter CD36 and exhibit increased β-oxidation and TCA cycle activity suggesting that LSCs likely fuel the TCA cycle and ETC with fatty acids from adipocytes (20–22). Recently, it has also been shown that AML cells, but not normal hematopoietic cells, provide free fatty acids to mitochondria via autophagy without exogenous supply (23). Independently of the fatty acid catabolism, enrichment of genes involved in lipogenesis has been reported within the purified subpopulation of human LSCs (24). These studies looking at lipolysis and lipogenesis support the idea that LSCs could have a different membrane composition from other leukemia cells and different from normal HSCs, which would explain the clear difference in cold sensitivity that we highlight here.

After isolation, thawing or for instance during flow cytometry staining procedures, cells are oftentimes kept at 4°C. We have observed that some samples can lose up to 50% viability in 6 hours. Additionally, we found that distinct subpopulations within primary samples, including LSCs, are even more vulnerable at 4°C than the bulk population. Beyond cell death characterized by permeabilization of the plasma membrane, it could also be that exposure to cold stimulates the differentiation of LSCs when they are returned to 37°C. Our data could explain why some primary patient samples have unexpectedly failed to engraft in immunodeficient mice following an overnight storage step at 4°C. If immediate use of the sample is not possible, the option of overnight storage at 4°C must be avoided and the samples should preferably be frozen.

Finally, although CKC4 cannot be applied to patients for direct therapeutic purposes, we believe that it could be tested in the context of autologous stem cell transplantation to selectively eradicate LSCs in order to avoid their reintroduction into the recipient patients. Current regulations do not provide specific information about the graft thawing method but underline that the thawing process must be carried out in the shortest possible time to bring the transplant to 37 °C. No incubation time at 4°C is used. Furthermore, our finding that OxPhos-driven AML cells and LSCs are sensitive to cold shock could possibly be extended to other blood malignancies and solid tumors and their respective Cancer Stem Cells.

## Acknowledgments

Authors are indebted to patients who granted permission to use samples in research. This work is dedicated to Dr Charles Pierre Griessinger. We thank the C3M core FCM Facility. EG thanks Ruxanda Moschoi for the first “failed experiment” which sparked the development of the CKC4. Part of this work was supported by an ERAPerMed-AML_PM grant to J.J.S and by grants from the Laboratoire d’Excellence Toulouse Cancer (TOUCAN and TOUCAN2.0; contract ANR11-LABEX) and the Ligue Nationale de Lute Contre le Cancer to J-E.S. JFP’s team projects are supported by INCa (PLBio2016-162), ANR (ANR-17-CE11-0002), Fondation ARC (PJA20191209626-1) and The Sohn Conference Foundation (Sohn Monaco). The Mediterranean Centre for Molecular Medicine is supported by institutional grants from INSERM. This study was supported by a grant from the Cancéropôle PACA and the Fondation de France (00067114).

## Author contributions

EG conceived the study and wrote the manuscript; EG, DPM, MN, CB, ES, EB, AS, JC, DD, LF, MP conducted experiments; JFP, JJS, JC, JES helped in conception and design of experiments; VD, FV, CR, GH, JES, provided cells and patients materials and data. EG, JFP, JJS, JES, DPM, JC reviewed and edited the manuscript.

## Methods

### Cell lines and culture conditions

Human AML cell lines were maintained in RPMI1640, with 10% fetal bovine serum (Lonza), 2mM L-glutamine, 1 mM sodium pyruvate, penicillin (50 U/ml) and streptomycin (50 mg/ml). TF-1 cells were supplemented with GM-CSF 10 ng/ml (Peprotech). AS-E2 cells were cultured in Iscove’s Modified Dulbecco’s Media (IMDM) supplemented with 20% fetal bovine serum and supplemented with EPO 5 U/ml. All cell lines were grown at 37 °C with 5% CO_2_, split every 2–4 days and maintained in an exponential growth phase. All AML cell lines were purchased at DSMZ or ATCC, and their liquid nitrogen stock was renewed every 2 years. These cell lines have been routinely tested for *Mycoplasma* contamination in the laboratory and monitored for metabolic drift.

### Patient Samples

Neonatal cord blood (CB) samples were obtained from healthy full-term pregnancies and mobilized peripheral blood from healthy donors were obtained at the obstetrics departments at the Martini Hospital and University Medical Center Groningen. Bone marrow or peripheral blood samples were obtained of patients diagnosed with de novo AML. All human samples, including healthy CB and AML samples, were obtained and studied after informed consent and protocol approval by the Ethical Committee in accordance with the Declaration of Helsinki.. CB CD34^+^ cells were isolated by density gradient separation, followed using a hematopoietic progenitor magnetic associated cell sorting kit (#130-046-703, Miltenyi Biotech) according to the manufacturer’s instructions. All patient tissue collection and research use adhered to protocols approved by the institutional review, in accordance with the Declaration of Helsinki. Details of patient samples are listed in supplementary Table 1. Mononuclear cells (MNCs) were isolated using LymphoPrep™ (Axis Shield Poc As™) separation and cryopreserved.

### Cold Killing Challenge at 4°C (CKC4)

Cells were incubated in a refrigerator calibrated at 4 degrees Celsius in RPMI 1640, with 10% fetal bovine serum, 2 mM L-glutamine, 1 mM pyruvate, penicillin (50 U/ml), streptomycin (50 mg/ml).

Temperature was controlled for each experiment with a precision gallium thermometer. After incubation cells were stained with DAPI (4,6 diamidino-2-phenylindole, Invitrogen) and analyzed at room temperature by Flow cytometry (FCM). For rotenone (R8875, Sigma-Aldrich), cells were treated for 10 minutes at 37°C, then washed in PBS and resuspend in a fresh medium and then incubated for the indicated time. Where indicated, glucose free RPMI medium (ThermoFisher) was supplemented with 5 mM galactose (Sigma-Aldrich). NB-4 cells were treated with 1 μM of retinoic acid (ATRA, Sigma-Aldrich) or control DMSO for the indicated time prior to CKC4. In some experiments, cells were treated for 48 to 72h with Etomoxir (#HY-50202, MedChem express) or Triacsin C (#ab141888, Abcam) with IC25 dose specific of each cell tested and predetermined by luciferase assay. After treatment cells were washed with PBS, then fresh medium was added prior to CKC4.

### Oxygen consumption measurement

Oxygen consumption was measured using Oxygen consumption rate (OCR) was measured in real time using the XF96 extracellular flux analyzer (Agilent). AML cells were seeded on Cell-Tak (#10317081, Thermo Fisher Scientific)-coated XF96 plates at 0.12×10^6^ cells/well in 180 μL of XF base medium minimal DMEM media (#102353-100, Agilent) supplemented with 20 mM D-glucose (G6152, Sigma-Aldrich), 1 mM sodium pyruvate (#11360088, Thermo Fisher Scientific), 2 mM L-glutamine (#25030081, Thermo Fisher Scientific) and adjusted to pH 7.4. The plates were spun at 200 g (breaks off) and incubated at 37°C for 20 min to ensure cell attachment. Measurements were obtained under basal conditions and in response to mitochondrial inhibitors, 1 μM oligomycin (O4876, Sigma-Aldrich), 0.5 μM of Carbonyl cyanide 4 (trifluoromethoxy) phenylhydrazone (FCCP, C2920, Sigma-Aldrich), and 1 μM rotenone (R8875, Sigma-Aldrich) combined with 2 μM antimycin A (A8674, Sigma-Aldrich).

### ATP Analysis

50,000 *luciferase* transduced cells were resuspended in 100µL of corresponding medium supplemented with 10% FBS and distributed in a 96-well plate. Cells were then treated in triplicates for two hours in glucose-free medium with galactose 2 g/L to inhibit glycolysis derived ATP, or in glucose medium with 2 µM of oligomycin to inhibit OxPhos derived ATP or in combination of both conditions to obtain the residual amount of ATP. Short time treatment (2 hours) was chosen to avoid a metabolic switch upon inhibition of glycolysis, to reach the maximum reduction effect and to avoid cell death upon inhibition of both metabolic pathways. D-Luciferin (#122799, Perkin Elmer) was added to give a final concentration of 150 μg/mL prior to luminescence measurement. Plates were analyzed with a Luminoscan (Berthold Technologies). We verified that ATP measurements were in the linear range of the detection. The difference between total ATP production (untreated condition) and the ATP produced under glucose-free medium with galactose treatment results in glycolytic ATP. The difference between total ATP production and the ATP produced under oligomycin treatment results in OxPhos ATP contribution. OxPhos and glycolytic derived ATP are represented by the percentage of total ATP produced by the cells.

### Flow Cytometry (FCM) Analysis

FCM Analyses were performed using MACSQuant, MACSQuant® VYB or MACSQuantX analysers (Miltenyi Biotec) or LSRFortessa or CytoFLEX flow cytometer. To distinguish the initial high cellular death inherent to the thawing of primary AML cells, mononuclear cells were pre-stained with the dead cell-specific amine-reactive fluorescent non-leaking dye Zombie Aqua™ (Supplementary Figure 4a). Human hematopoietic cells were first stained with Zombie Aqua™ (#423101, Biolegend) following the manufacturer’s instructions. After washing cells were stained with CD45-APC-Cy7 (#304014, Biolegend), CD34-PE (#345802, BD Biosciences), CD38-APC (#303510 Biolegend), CD123-PECy7 (#560826 BD Biosciences), CD45RA-FITC (#304106 Biolegend) antibodies. DAPI was added at room temperature prior to FCM analysis. The negative fraction was determined using appropriate isotype antibodies and live population control. For measuring mitochondrial potential at 37°C or mitochondrial permeability at 4°C, cells were stained with 1 μM of Tetramethylrhodamine, ethyl ester, perchlorate (#87917, TMRE, Sigma-Aldrich) for 30 minutes at 37°C. For FCM analyses were performed with FlowJo software (TreeStar). HSC, LMPP, MEP, CMP, and GMP-like gates enclosing less than 300 events were excluded for viability analysis.

### Mass Spectrometry data acquisition

Samples were analyzed by direct infusion on a QExactive mass spectrometer (Thermo Scientific) equipped with a TriVersa NanoMate ion source (Advion Biosciences). Samples were analyzed in both positive and negative ion modes with a resolution of Rm/z=200=280000 for MS and Rm/z=200=17500 for MS/MS experiments, in a single acquisition. MS/MS was triggered by an inclusion list encompassing corresponding MS mass ranges scanned in 1 Da increments. Both MS and MS/MS data were combined to monitor CE, DAG and TAG ions as ammonium adducts; PC, PC O-, as acetate adducts; and CL, PA, PE, PE O-, PG, PI, and PS as deprotonated anions. MS only was used to monitor LPA, LPE, LPE O-, LPI, and LPS as deprotonated anions; Cer, HexCer, SM, LPC and LPC O-as acetate adducts.

### Long Term Culture (LTC)

Co-culture experiments were performed as previously described (15) as bulk culture or using a limiting dilution analysis (LDA) both on a confluent monolayer of MS-5, supplemented with recombinant human IL3, G-CSF, and TPO (MS-5 + 3GT) (20 ng/mL each; Peprotech) in Gartner’s medium consisting of αMEM (Thermo Scientific) supplemented with 12.5% heat-inactivated fetal bovine serum (Lonza), 12.5% heat-inactivated horse serum (Invitrogen), 1% penicillin and streptomycin, 2 mM glutamine (all from PAA Laboratories).

### Xenograft assay

NOD-SCID common γ-chain knockout mice (NSG mice) were purchased from the Centrale Dienst Proefdieren (CDP) breeding facility within the University Medical Center Groningen. Mouse experiments were performed in accordance with national and institutional guidelines, and all experiments were approved by the Institutional Animal Care and Use Committee of the University of Groningen (IACUC-RuG). Healthy 6- to 8-wk-old female mice received a single i.v. injection of 1.4× 10^6^ DAPI negative primary AML cells (sample #35) at T0 or 24 h post CKC4. Animals were sacrificed 10 weeks after injection and cells from mouse PB, BM and Spl were isolated. Leukocytes were recovered after red cells lysis with ammonium chloride. Cells were stained with human specific APC– conjugated anti-CD19, PE-conjugated anti-CD33, PE-Cy7–conjugated anti-CD45 (all from BD Pharmingen). AML engraftment was defined by the presence of a single CD45^+^CD33^+^CD19^-^ population.

### Statistics

Data presented is either a mean of triplicates samples from one representative experiment reproduced independently or a Grand Mean of Mean response of up to 11 merged independent experiments themselves made in triplicates. Data were analyzed for statistical significance using the Mann-Whitney unpaired two tail test or the one-way ANOVA test. A non-parametric Spearman test was applied for correlation. Spearman’s rank correlation coefficient (ρ) is shown. Observed differences were regarded as statistically significant if the calculated two-sided P value was below 0.05. When a correlation was proven a linear or non-linear regression trend lines were performed with GraphPad Prism software (v9.5.0).

