## Supplementary text and figures for "FAO-supported OxPhos leukemic stem cells are sensitive to cold"

***List of Supplementary data*:**

***Supplementary method.***
***Supplementary Figure 1. OxPhos Leukemic Cells Exhibit a specific and reproducible sensitivity to 4°C and resulting in a faster permeabilization of their membrane during CKC4 compared to glycolytic cells*.**
***Supplementary Figure 2. OxPhos leukemic cells have a CKC4 sensitive specific lipidome dependent on fatty acid metabolism.***
***Supplementary references.***

***Table S1:*** *Characteristics of AML patient samples.*

Supplementary data:

#### Supplementary method.

Lactate production

Lactate concentration in the supernatant was determined electro-enzymatically using the YSI 2950 Biochemistry Analyzer (Yellow Springs Instruments). Lactate secretion was normalized to viable cell numbers (using Trypan blue exclusion method) and expressed as picograms per cell per min. Alternatively lactate was expressed as gram per liter when the same cellular density was verified in the different conditions. Extracellular lactate concentration was also determined by the lactate dehydrogenase enzymatic reaction in the cell culture medium at 0, 24, and 48 h of incubation. Extracellular lactate was converted by L-Lactic Dehydrogenase (LDH, Sigma Aldrich) reaction in freshly prepared 25 mM $NAD^+$ and 87.7 U/mL LDH in 0.4 M hydrazine (Sigma Aldrich)/0.5 M glycine assay buffer (pH 9). 20 µL samples (diluted according to standard curve) and Sodium L-lactate (Sigma Aldrich) standards were pipetted into 130 µL reagent mix in 96 wells plate format and the reaction was carried out for 30 min at 37 °C. Measurements were recorded from a microplate reader at 340 nm wavelength (Multiskan Sky microplate reader, Thermo Fisher Scientific).

Lipid extraction for mass spectrometry lipidomics

Mass spectrometry-based lipid analysis was performed by Lipotype GmbH (Dresden, Germany) as described (1). Lipids were extracted using a two-step chloroform/methanol procedure (2). Samples were spiked with internal lipid standard mixture containing: cardiolipin 16:1/15:0/15:0/15:0 (CL), ceramide 18:1;2/17:0 (Cer), diacylglycerol 17:0/17:0 (DAG), hexosylceramide 18:1;2/12:0 (HexCer), lyso-phosphatidate 17:0 (LPA), lyso-phosphatidylcholine 12:0 (LPC), lyso-phosphatidylethanolamine 17:1 (LPE), lyso-phosphatidylglycerol 17:1 (LPG), lyso-phosphatidylinositol 17:1 (LPI), lyso-phosphatidylserine 17:1 (LPS), phosphatidate 17:0/17:0 (PA), phosphatidylcholine 17:0/17:0 (PC), phosphatidylethanolamine 17:0/17:0 (PE), phosphatidylglycerol 17:0/17:0 (PG), phosphatidylinositol 16:0/16:0 (PI), phosphatidylserine 17:0/17:0 (PS), cholesterol ester 20:0 (CE), sphingomyelin 18:1;2/12:0;0 (SM), triacylglycerol 17:0/17:0/17:0 (TAG). After extraction, the organic phase was transferred to an infusion plate and dried in a speed vacuum concentrator. 1st step dry extract was re-suspended in 7.5 mM ammonium acetate in chloroform/methanol/propanol (1:2:4, V:V:V) and 2nd step dry extract in 33% ethanol solution of methylamine in chloroform/methanol (0.003:5:1; V:V:V). All liquid handling steps were performed using Hamilton Robotics STARlet robotic platform with the Anti Droplet Control feature for organic solvents pipetting.

Mass Spectrometry data analysis and post-processing.

Data were analyzed with in-house developed lipid identification software based on LipidXplorer (3,4). Data post-processing and normalization were performed using an in-house developed data management system. Only lipid identifications with a signal-to-noise ratio >5, and a signal intensity 5-fold higher than in corresponding blank samples were considered for further data analysis. Each sample was analyzed in triplicate. Lipid species mol% values per sample were used as input data for the PCA score plot.

Proteome and Pathway Analysis

Proteome details regarding detection and quantification of the full label-free quantitative proteome are outlined in our previously published study (5). Here 10 AML patient samples have been used. Pearson's correlation value was determined between Q1V$^-$ time parameter and the proteome data. The protein list from the cell line CCLE metabolomic data and from our patient cohort was selected based on a Pearson's correlation value (rho) > 0.3. Gene Ontology (GO)-analysis was performed using the online geneontology.org (6). Gene set enrichment analysis (GSEA, v4.1.0) was performed on a pre-ranked gene list with respect to MSigDB genesets (version 7.3)(7).

Mitochondrial DNA content (mtDNAc)

Total DNA was isolated from pelleted cells (0.5-2x$10^6$ cells) with the NucleoSpin® Tissue kit according to the manufacturer's protocol. DNA concentration was measured with a Nanodrop™ spectrophotometer (Thermo Scientific) and 10 ng of DNA was used for each reaction. mtDNA copy number was performed according to Panero et al. (8) by quantitative RT-PCR. Briefly, *MT-ND4* and *MT-CYB* genes were used to represent the mtDNA, and pyruvate kinase (*PK*) gene and the beta-globin (*HBB*) gene were used to represent the nuclear DNA. Relative mtDNA copy numbers were assessed after *MT-ND4*/*CYB* normalization by the average expression for the single-copy nuclear gene *PK* and *HBB*. All reactions were performed using SsoAdvanced SYBR Green Supermix (Bio-Rad) on a CFX384 Touch Real-Time PCR Detection System (Bio-Rad). The corresponding real-time PCR efficiencies for each mitochondrial and nuclear gene amplification were calculated according to the equation: $E = 10(-1/slope) - 1$. The relative mtDNA copy number was calculated relative to a reference DNA sample (healthy donor) and set to 1. In all experiments, the same reference DNA was used as an internal control to ensure that the results would be fully comparable among experiments. The relative mtDNA copy number was calculated as relative quantification using the ΔCt method and expressing the results as $2^{-\Delta\Delta Ct}$, in which $\Delta\Delta Ct = \Delta Ct_{sample} - \Delta Ct_{reference\ DNA}$.

*Long Term Culture (LTC)*

For LTC in limiting dilution analysis, cells were plated in 20 replicates in 96-well microplates containing confluent MS-5 monolayer. The murine bone marrow stromal cell line MS-5 was kindly provided by Dr Bonnet (The Francis Crick Institute, London, UK) and maintained in IMDM 10% FBS plus 2 mM L-glutamine. Half medium change was done twice a week without disrupting the established feeders. After 5 weeks, the LTC medium was replaced by methylcellulose (#H4435, StemCell Technologies). After an additional 2 weeks, each well was scored as negative if no colonies were present. The frequency of normal and leukemic LTC-IC was determined using ELDA (WEHI, Bioinformatics Division) (9) according to the Poisson statistics and method of maximum likelihood.

### Supplemental Figure 1

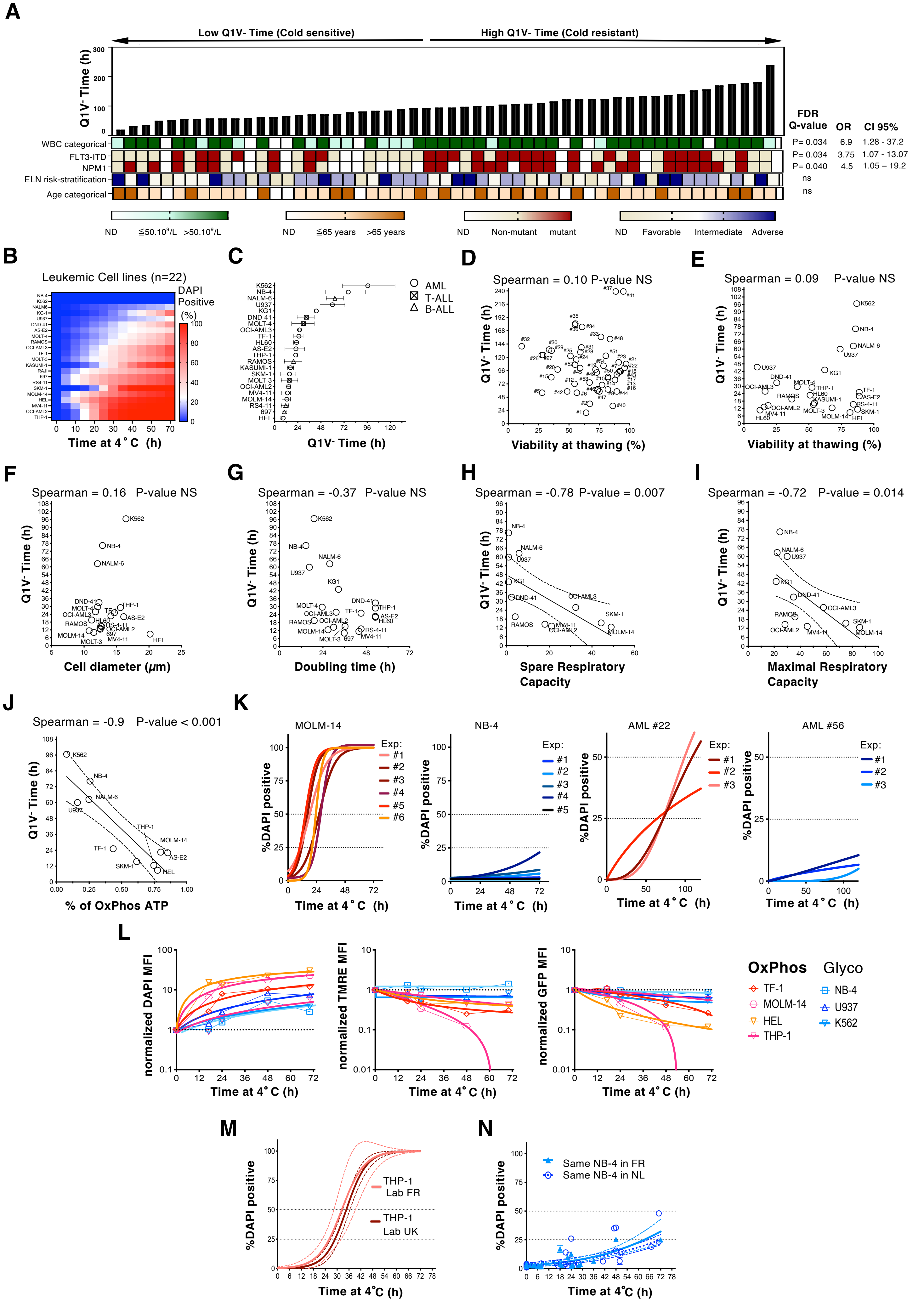

**Figure S1. Related to Figure 1. OxPhos Leukemic Cells Exhibit a specific and reproducible sensitivity to 4°C and resulting in a faster permeabilization of their membrane during CKC4 compared to glycolytic cells.** (A) Distribution of clinical characteristic and genetic abnormalities in AML with low and high Q1V$^-$ Time. Discrete state/class is indicated as colour. White Blood Cell Count (WBC), *FLT3* and *NPM1* genetic alterations, European LeukemiaNet (ELN) recommendations, age. Significance was determined using either two-tailed Mann–Whitney or Kruskal–Wallis tests (for categorical variables). Only mutations that were found in ≥3 patients were considered. Significance was determined using a Kruskal–Wallis test. (B) Heatmap showing the kinetics of mean viability for 21 leukemic cell lines submitted to a cold killing challenge at 4°C (CKC4). For each cell line a non-linear regression trend line was derived from 3 to 15 independent CKC4 experiments. (C) Time to lose the first quartile of viability at 4°C (Q1V$^-$ Time) (± SEM) of AML, T-ALL and B-ALL cell lines. (D-E) Q1V$^-$ Time against the cell viability after thawing from -196°C for primary AML samples (n=48 pairs) and leukemic cell lines (n=18 pairs). Viability after thawing is expressed as mean values derived from 2 to 3 independent thawing experiments. (F) Q1V$^-$ Time against the cell diameter for indicated leukemic cell lines (n=18 pairs). (G) Q1V$^-$ Time against the cell doubling time at 37°C. The mean doubling time was determined by cell count performed in exponential growth. (H-I) Spare and Maximal Respiratory Capacity as a function of Q1V$^-$ Time. For each cell line, metabolites measurements were performed 2-6 times in independent experiments or at different time points. (J) Mitochondrial ATP production rate as compared to Q1V$^-$ Time. Luciferase-expressing AML cell lines were treated with oligomycin A or incubated in a glucose free medium with galactose to respectively block oxidative phosphorylation or glycolysis, or a combination of both conditions. The glycolytic and mitochondrial ATP production rate was measured using the luciferase assay. For D-J a nonparametric Spearman correlation test was applied. Thin black line shows experimental derived linear regression trend line with 95% confidence band. (K) Viability kinetics of MOLM-14, NB-4, primary AML #22 and #56 cells determined in independent CKC4 experiments. (L) Fluorescence leak in and leak out for four OxPhos and three glycolytic leukemic cells during CKC4. Data show the kinetics of variation of median fluorescence intensity by FACS of DAPI (DAPIMFI, left), TMRE MFI (middle), GFP MFI (right) during CKC4. For each cell line the MFI was normalized by the initial MFI at T0 of the considered fluorescence. DAPI (inward leaking dye, added extemporaneously of the CKC4) fluorescence increases overtime. GFP (outward leaking fluorescent protein, constitutively expressed by transduction prior to the CKC4) and TMRE (outward leaking dye, stained cells prior to the CKC4) fluorescence decrease overtime. Doublets and debris were eliminated by forward and side light scatter. (M) CKC4 kinetic profiling of THP-1 cells issued from the laboratories cell collection of the HSCL-lab (gift from Pr D. Bonnet; London, UK) and of the TrGET Preclinical Platform (gift from Y. Collette; Marseille, FR). (N) CKC4 kinetic profiling of the same batch of NB-4 cells performed at C3M institute (INSERM U1065, Nice, FR) and at UMCG (Groningen, NL) with an interval of one year. Thin line shows experimental derived non-linear regression trend line with 95% confidence band (dashed lines).

### Supplemental Figure 2

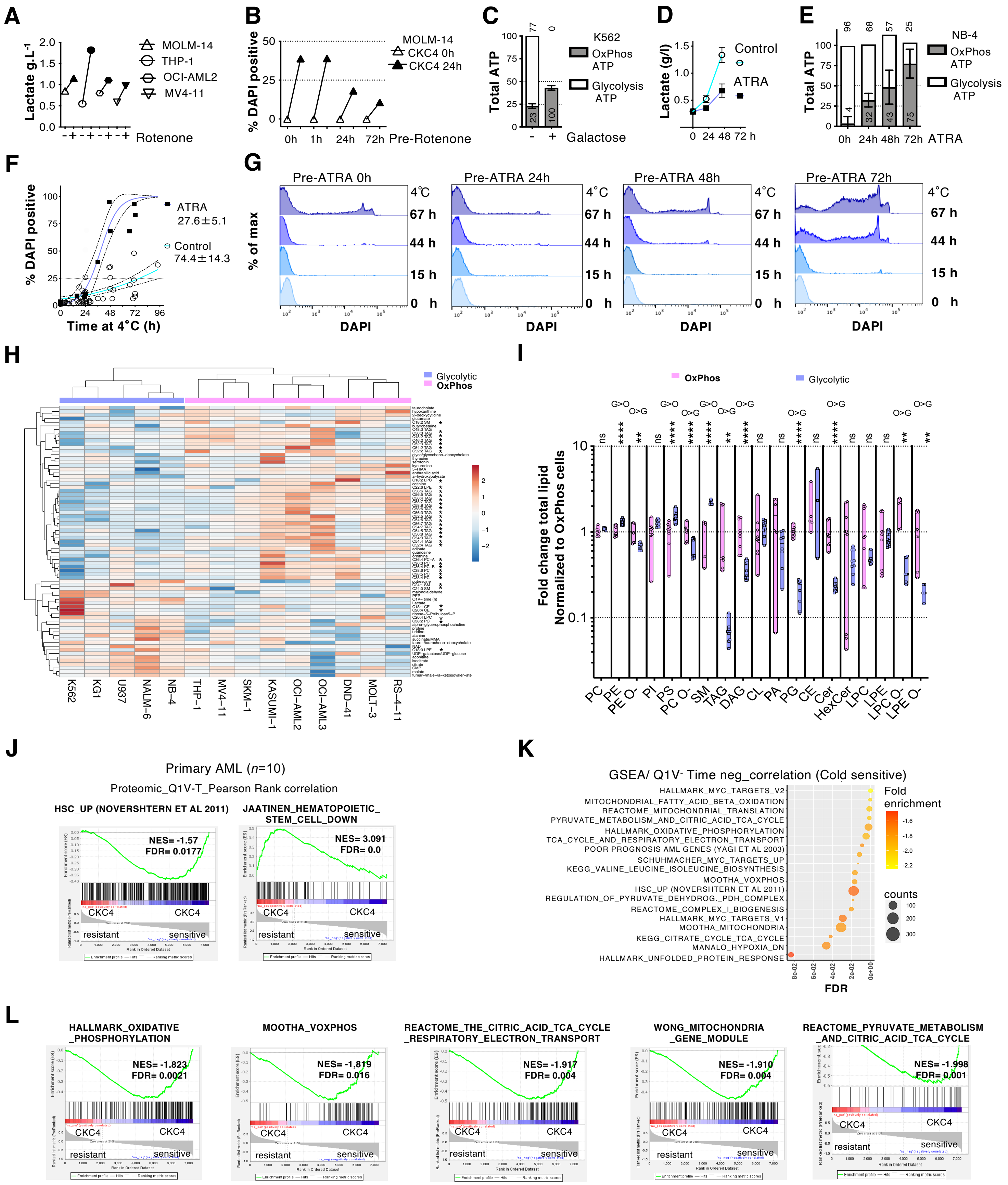

**Figure S2. Related to Figure 2. OxPhos leukemic cells have a CKC4 sensitive specific lipidome dependent on fatty acid metabolism.** (A-B) Shift of OxPhos cells toward Glycolytic metabolism. Mitochondrial respiration was impaired by the complex I inhibitor rotenone. (A) Mitochondrial respiration of MOLM-14 THP-1 MV4-11 and OCI-AML2 cells was impaired by the irreversible complex I inhibitor rotenone. After 10 min treatment with rotenone 0.1 µM, cells were washed and incubated at 37°C for 72h and lactate was measured in culture medium for the same output number of treated or untreated control cells. (B) Rotenone pre-treated (Pre-rotenone) MOLM-14 cells were incubated at 37°C for 1 or 24 or 72h before undergoing a CKC4 for 24h. Data show the progressive decline of DAPI positivity after CKC4 for increasing time of metabolic reprogramming. (C-G) Shift of glycolytic cells toward OxPhos metabolism. (C) Glycolytic and mitochondrial ATP production rate (respective percentages displayed) of K562 cells incubated at 37°C in glucose-free medium with galactose for 72h measured by luciferase assay. Total ATP was normalized to untreated control. (D) Lactate concentration overtime in cell culture media at 37°C of ATRA 1 µM 72h treated versus control NB-4. To compensate for cytostatic effect of ATRA, a larger number of cells were incubated under ATRA condition in order to measure lactate production for an equivalent number of cells. (E) Glycolytic and mitochondrial ATP production rate normalized to untreated control of NB-4 cells treated with ATRA for 24, 48 or 72h at 37°C. (F) CKC4 kinetic profiling of ATRA treated versus control NB-4 cells. Thin black line shows experimental derived linear regression trend line with 95% confidence band (dashed lines). The data was combined from three independent experiments. The derived $Q1V^-$ Time (± SEM) of ATRA treated versus control NB-4 cells is displayed. (G) NB-4 cells were incubated with ATRA at 37°C for 24, 48 or 72h before undergoing a CKC4 for 15, 44 or 67h. FACS histogram plot showing DAPI incorporation. Data show the progressive increase of DAPI positivity after CKC4 for increasing time of metabolic reprogramming. (H) Heat map showing the hierarchical clustering of CCLE metabolomic data for the top 70 metabolites that were significantly different between OxPhos and glycolytic cells and correlating with the $Q1V^-$ Time parameter. Stars indicate lipid among the metabolite's entries. (I) Fold change of total lipid per class for OxPhos (O: Kasumi-1, THP-1, MV4-11) and glycolytic cells (G: KG-1a, NB-4 and U937) normalized to the mean percentage for OxPhos cells (see also Table S1 and Fig. S3). Lipidome was measured in triplicate per cell type by high-resolution Orbitrap mass spectrometry. A Mann–Whitney test was applied, ns $p>0.05$, $^{**}p < 0.01$, $^{****}p < 0.0001$. (J-L) Total proteome by MS for 10 patient samples. Pearson correlations were calculated for all quantified proteins versus $Q1V^-$ Time determined, and a ranked list of Pearson coefficients was used to perform gene set enrichment analyses. (J) GSEA analysis of HSC up and HSC down pathways in CKC4 resistant versus CKC4 sensitive primary AML samples. (K) Pathway enrichment bubble plot of differentially expressed genes negatively correlated with $Q1V^-$ Time (Cold sensitivity). The number of associated genes in the pathway is indicated by the circle area, and the circle colour represents the range of the correlation. (L) GSEA analysis of oxidative phosphorylation Oxidative phosphorylation, TCA cycle respiratory electron transport, mitochondrial gene module and Pyruvate metabolism and Citric Acid (TCA) cycle pathways in CKC4 resistant versus CKC4 sensitive primary AML samples.

**Supplementary References.**
